# Genome-wide evidence of the role of Paf1C in transcription elongation and histone H2B monoubiquitination in *Arabidopsis*

**DOI:** 10.1101/2023.10.27.564443

**Authors:** Noel Blanco-Touriñán, Jaime Pérez-Alemany, Clara Bourbousse, David Latrasse, Ouardia Ait-Mohamed, Moussa Benhamed, Fredy Barneche, Miguel A. Blázquez, Javier Gallego-Bartolomé, David Alabadí

## Abstract

The evolutionarily conserved Paf1 complex (Paf1C) participates in transcription, and research in animals and fungi suggests that it facilitates RNAPII progression through chromatin remodeling. To obtain evidence that Paf1C acts in transcription elongation in *Arabidopsis*, we examined the genomic distribution of the ELF7 and VIP3 subunits of Paf1C. The occupancy of both subunits was confined to thousands of gene bodies and positively correlated with RNAPII occupancy and the level of gene expression, supporting a role as a transcription elongation factor. We found that monoubiquitinated histone H2B, which marks most transcribed genes, was strongly reduced genome-wide in *elf7* seedlings. Genome-wide profiling of RNAPII revealed that in *elf7* mutants, RNAPII occupancy was reduced throughout the gene body and at the transcription end site of Paf1C-targeted genes, suggesting a direct role for the complex in transcription elongation. Overall, our observations suggest that there is a direct functional link between Paf1C activity, mono-ubiquitination of histone H2B, and the transition of RNPII to productive elongation. However, for several genes, Paf1C may also act independently of H2Bub deposition or occupy these genes more stably than H2Bub marking, possibly reflecting the dynamic nature of Paf1C association and H2Bub turnover during transcription.

## INTRODUCTION

During gene transcription, numerous proteins and protein complexes help RNA Polymerase II (RNAPII) to move through the gene body, including histone chaperones, chromatin remodelers, and transcription elongation factors, most of which are evolutionarily conserved in eukaryotes (Obermeyer et al., 2023). One of them is the transcription elongation complex POLYMERASE ASSOCIATED FACTOR1 (Paf1C). Paf1C was originally identified in yeast and the current view of how Paf1C acts in gene regulation is largely based on studies in yeast and mammals (Francette et al., 2021). Early genetic and biochemical evidence for a positive role of Paf1C in transcriptional elongation (Shi et al., 1997; Rondón et al., 2004) is now supported by structural studies showing the association of the complex with active RNAPII (Vos et al., 2018; Vos et al., 2020). Paf1C is also involved in posttranslational modification of histones. Histone modifications such as the monoubiquitination of histone H2B (H2Bub) or the methylation of lysines 4 or 36 of histone H3 (H3K4me, H3K36me), which influence chromatin dynamics co-transcriptionally to facilitate RNAPII progression, have been found to be dependent on Paf1C in yeast and mammals (Francette et al., 2021).

Several subunits of plant Paf1C have been genetically identified in *Arabidopsis* in genetic screens for early flowering mutants or by searching for homologous genes (Zhang and Van Nocker, 2002; Zhang et al., 2003; He et al., 2004; Oh et al., 2004; Park et al., 2010; Yu and Michaels, 2010). As in animals, the plant Paf1C contains six subunits, PLANT HOMOLOGOUS TO PARAFIBROMIN (PHP)/CELL DIVISION CYCLE73 (CDC73), VERNALIZATION INDEPENDENCE2 (VIP2; also known as EARLY FLOWERING7, ELF7), VIP3, VIP4, VIP5, and VIP6 (ELF8), which are the putative orthologs of CDC73, Paf1, SKI8, LEO1, RTF1, and CTR9, respectively (Obermeyer et al., 2023). The *Arabidopsis* Paf1C is formed *in vivo*, since at least VIP3, VIP4, and VIP6 interact with PHP/CDC73 (Park et al., 2010) and all subunits were identified in interactomic analyses using CDC73 and ELF7 as baits (Antosz et al., 2017). Furthermore, the latter study showed that Paf1C interacts with *bona fide* transcriptional elongation factors and with several subunits of RNAPII, pointing to a conserved role in transcriptional elongation.

In addition to flowering, and consistent with the pleiotropic effects of loss-of-function mutants in several subunits, plant Paf1C has been associated genetically with a variety of largely unrelated physiological processes in *Arabidopsis*, including seed dormancy (Liu et al., 2011), shoot apical meristem activity (Fal et al., 2017; Fal et al., 2019; Li et al., 2022), thermomorphogenesis (Zhao et al., 2023), and the response to salt stress (Zhang et al., 2022) and DNA damaging agents (Li et al., 2023).

In *Arabidopsis*, the role of Paf1C in post-translational modification of histones is well-documented and has been associated with several physiological processes. This includes the regulation of flowering time by acting on the flowering repressor gene *FLOWERING LOCUS C* (*FLC*) and its paralogs (Cao et al., 2015; Lu et al., 2017; Li et al., 2019; Nasim et al., 2022). The requirement of Paf1C for histone modifications has also been shown at the epigenomic level. Although the total amount of H3K4me3 and H3K36me2 marks is not altered in *vip3*, *vip4*, *vip5*, or *vip6* cell extracts (Oh et al., 2004), the *vip3* mutation triggers a slight redistribution of both marks across gene bodies (Oh et al., 2008). Moreover, plant Paf1 subunits are required for the correct distribution of H2A.Z and H3.3 in response to warm temperatures (Zhao et al., 2023) and for regulating H3K27 trimethylation at a subset of genes (Oh et al., 2008; Schmitz et al., 2009).

In line with the role of Paf1C in animals and fugi in promoting H2Bub accumulation, mutations in *Arabidopsis VIP2, VIP4, VIP5,* and *VIP6* subunits cause decreased H2Bub accumulation at *FLC* and its paralog genes (Cao et al., 2015). More recently, the dependence of H2Bub on Paf1C has been linked to the DNA damage response (Li et al., 2023). In *Arabidopsis*, as in yeast and animals, H2Bub accumulates across the gene bodies of actively transcribed genes (Roudier et al., 2011; Bourbousse et al., 2012; Nassrallah et al., 2018). However, it is not known whether Paf1C is present at many transcribed genes where it might contribute to H2Bub deposition or whether it indifferently affects this mark at all *loci*. To gain insight into transcription-associated chromatin mechanisms and their regulatory roles in plant development and stress responses, we here examined the involvement of Paf1C in transcription elongation by RNAPII and in monoubiquitination of histone H2Bub throughout the *Arabidopsis* genome.

## RESULTS

### The subunits of Paf1C ELF7 and VIP3 are recruited to gene bodies

First, we investigated the genomic distribution of Paf1C by profiling the chromatin association of the ELF7 and VIP3 subunits by ChIP-seq in one-week-old *Arabidopsis* seedlings. ChIP-seq analyses were performed using the *Flag-ELF7* and *GFP-VIP3* transgenic lines, respectively (Dorcey et al., 2012; Cao et al., 2015) and non-transgenic WT plants as negative controls. The resulting analysis indicated a high correlation between biological replicates (Fig. S1, A and B) and identified 9,791 and 6,845 Flag-ELF7 and GFP-VIP3 peaks, respectively (usually one peak per protein-coding gene; see examples in Fig. S1C), of which the vast majority was common (Fig. 1, A and B and Supplemental Table). A heatmap representation of Flag-ELF7 and GFP-VIP3 ChIP signal over unique and shared targeted genes confirmed that most of them were in fact bound by both proteins, albeit with different intensities (Fig. S2A). Thus, for the next analyses we considered all the peaks identified for the two subunits as *bona fide* Paf1C target loci (10,579) (Fig. 1A).

**Figure 1.**
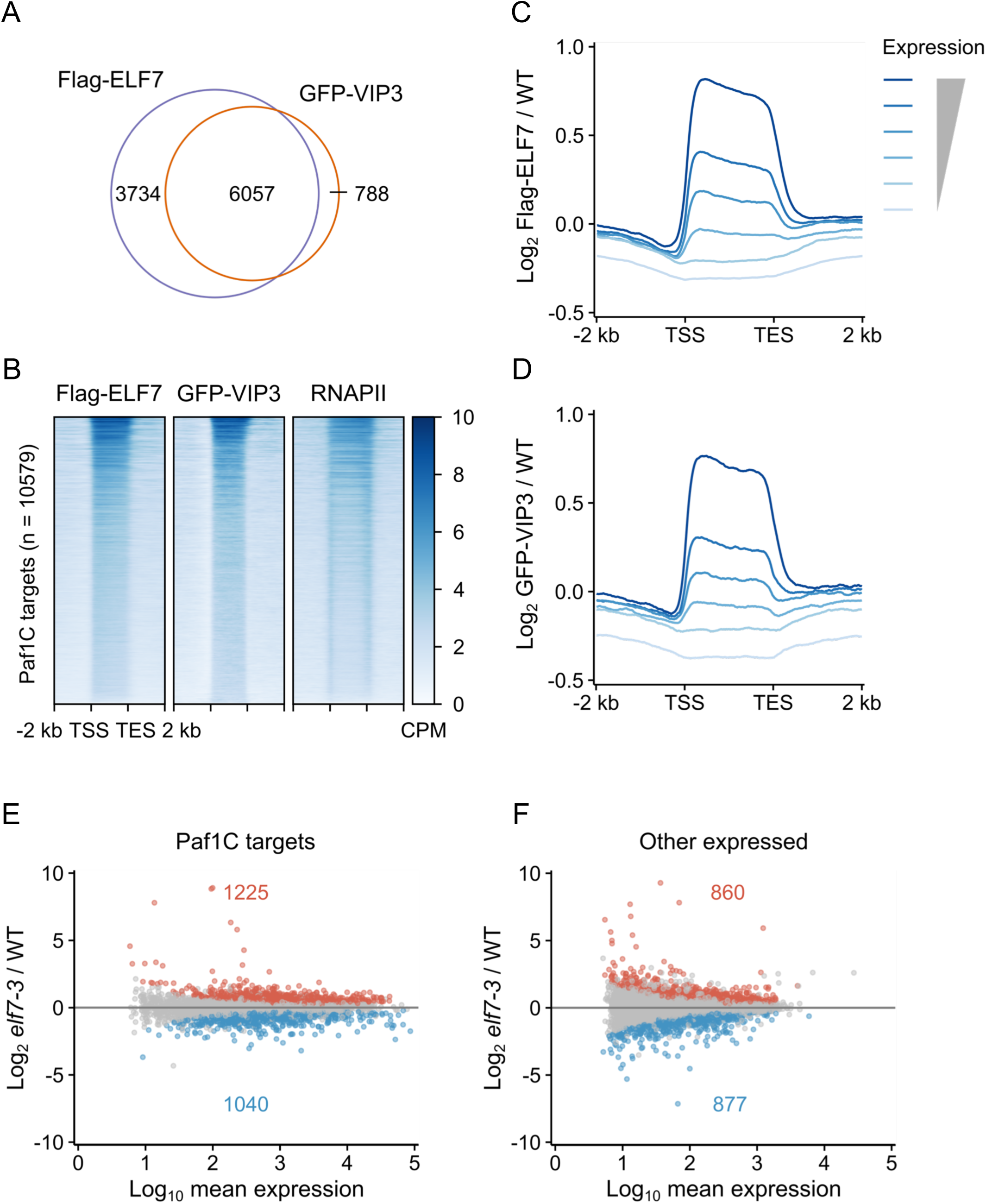
ELF7 and VIP3 bind to the gene body of transcribed genes. **A)** Venn diagram showing the overlap between Flag-ELF7 and GFP-VIP3 peaks. **B)** The heatmap shows the occupancy of Flag-ELF7, GFP-VIP3, and total RNAPII ranked by the decreasing Flag-ELF7 level. **C, D)** Average occupancy of Flag-ELF7 (C) and GFP-VIP3 (D) on target genes ranked by their expression level. **E, F)** MA plot with DEGs (*p* adj. < 0.05) in the *elf7-3* mutant compared with wild type for Paf1C target genes (E) and for genes in which we could not detect Paf1C (F).

The genomic profile of Paf1C showed a strongest enrichment over gene bodies that slightly extends in 3’ over the transcription end site (TES) (Fig. 1, B-D, and Fig. S2A). Comparison with RNAPII profiles showed that the overlap of the distributions of Paf1C and RNAPII was restricted to gene bodies (Fig. 1B). The ELF7 distribution observed in this work was very similar to that found in a recent study (Fig. S2B) (Wang et al., 2023b). In accordance with Paf1C’s role in transcription elongation, the occupancy of ELF7 and VIP3 exhibited a positive correlation with RNAPII occupancy (Fig. S1B) and transcript levels (Fig. 1, C and D). To further substantiate this perspective, we examined the accumulation of different histone marks and histone variants at Paf1C target genes (Jamge et al., 2023). Paf1C target genes were enriched in histone marks associated with active gene transcription, such as H3K4me3 and H3K36me3, while being depleted of the *Polycomb*-based H3K27me3 mark (Fig. S2A). Also consistent with Paf1C being associated with expressed genes and not with *Polycomb*-repressed genes, histone variant H2A.Z was enriched at the 5’ end but mostly absent across the gene bodies of Paf1C targets (Fig. S2A) (Coleman-Derr and Zilberman, 2012; Gómez-Zambrano et al., 2019).

To investigate the impact of Paf1C depletion on gene expression, we determined the transcriptome in wild-type and *elf7-3* one-week-old seedlings by RNA-seq. We identified 4,002 differentially expressed genes (DEGs; *p* adj. < 0.05) in *elf7-3* seedlings (Fig. 1, E and F, and Supplemental Table). More than half of the identified DEGs (2,265) were *bona fide* Paf1C targets (Fig. 1E and Fig. S2C). Interestingly, when we plotted the average signal of Flag-ELF7 or GFP-VIP3 across DEGs in the “Other expressed” category (Fig. 1F), we found enrichment in gene bodies for both subunits compared with gene bodies of non-expressed genes, which correlates with the RNAPII signal (Fig. S2C). This result suggests that the DEGs of the “Other expressed” category are also target genes of Paf1C, although they are only weakly bound by the complex in accordance with their low expression level. The weak binding may be the reason that these genes were below the threshold to be detected by peak calling as true Paf1C targets. In any case, the DEGs represent a fraction of all loci bound to Paf1C in the genome, a situation similar to that observed in human cells knocked-down for the ortholog Paf1 subunit (Yu et al., 2015). Among the *bona fide* Paf1C target DEGs, 1,040 were downregulated consistent with a positive role of the complex on transcription elongation. Intriguingly, 1,225 DEGs were upregulated in the mutant, suggesting a negative impact of Paf1C on their expression or the existence of indirect effects mediating their induction in the absence of Paf1C.

### Defects in the Paf1C complex affect H2Bub mark genome-wide

Paf1C is required for efficient monoubiquitination of H2B in various organisms, including plants (Wood et al., 2003; Schmitz et al., 2009; Hou et al., 2019; Li et al., 2023). To determine whether reduction of H2Bub deposition is detected at specific genes or on a global scale along the genome upon Paf1C loss-of-function, we examined the H2Bub dependence in wild-type and *elf7-3* seedlings by ChIP-seq with spike-in of exogenous chromatin (ChIP-Rx). Following the protocol described in Nassrallah et al. (2018) for *Arabidopsis*, spike-in based normalization enables for accurate quantitative comparisons of samples with genome-wide differences in chromatin mark abundances In our analyses, H2Bub was exclusively enriched in the transcribed regions (Fig. 2, A and B), in agreement with previous reports in *Arabidopsis* (Roudier et al., 2011; Bourbousse et al., 2012; Nassrallah et al., 2018). In addition, H2Bub-occupied genes were also co-ocupied by Paf1C and RNAPII (Fig. S3A), as expected for a co-transcriptionally deposited mark.

**Figure 2.**
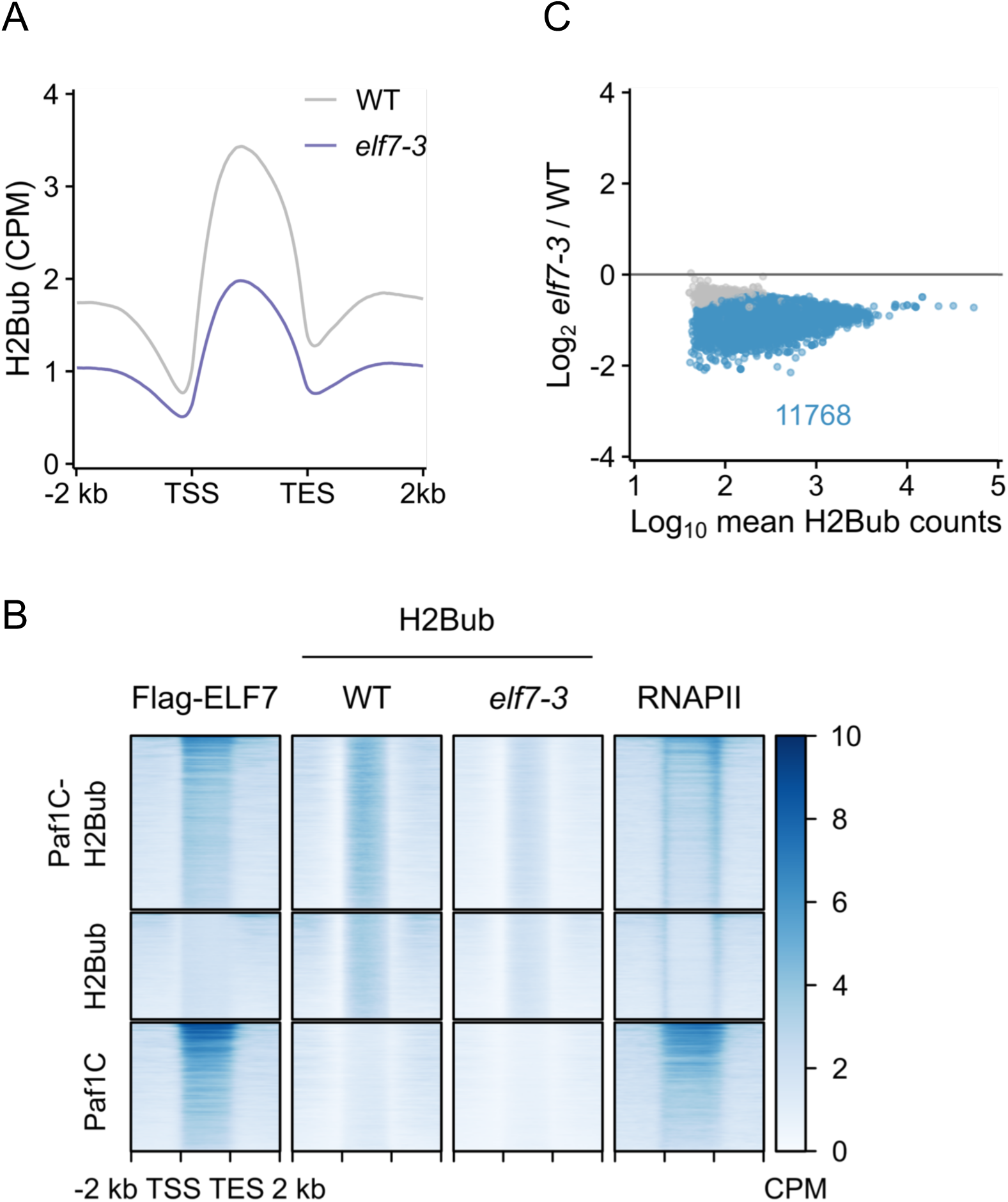
The H2Bub mark is reduced in *elf7-3* mutants.**A)** Average occupancy of the H2Bub mark in the wild type and in the *elf7-3* mutant. **B)** Heatmaps showing Flag-ELF7, H2Bub -in wild-type and *elf7-3* seedlings- and RNAPII occupancy in gene clusters based on H2Bub and Paf1C, H2Bub, or Paf1C occupancy. Genes are ranked by their decreasing levels of H2Bub in wild type. **C)** The MA plot shows differentially ubiquitinated genes (blue dots; *p* adj. < 0.05) between the wild type and the *elf7-3* mutant.

In the absence of Paf1C, H2Bub levels were strongly affected at multiple genes (Fig. 2, A and B, and Fig. S3, B and C). More precisely, identification of differentially ubiquitinated genes showed a general tendency for reduced H2Bub enrichment over most H2Bub-marked genes in *elf7-3* seedlings (Fig. 2C and Supplemental Table). Hence, given the frequent local co-occurrence of Flag-ELF7/GFP-VIP3 and the H2Bub mark, these observations suggest the existence of direct functional links between Paf1c activity and histone H2B monoubiquitination.

However, the association of Flag-ELF7 and GFP-VIP3 with chromatin was also found in genes enriched in RNAPII but not H2Bub (Fig. 2B, and Fig. S3C and Supplemental Table). We found that these genes were smaller compared with the genes that were co-ocuppied by Paf1C and H2Bub (Fig. S3D). As previously identified using transcriptome data (Bourbousse et al., 2012), this observation suggests that optimal transcription of small genes in *Arabidopsis* is less dependent on H2Bub. Furthermore, it indicates that Paf1C may function independently of H2Bub deposition at multiple genes or that it may occupy these genes more persistently than H2Bub. *Vice versa*, H2Bub enrichment was detected at genes in which we could not detect Paf1C (Fig. 2B). Interestingly, H2Bub was also reduced at these genes in *elf7-3* seedlings, possibly indicating that Paf1C acted on histone H2B monoubiquitination transiently, or earlier, at these loci. In summary, these observations suggest that histone H2B monoubiquitination is tightly linked to Paf1C activity at many genes but may also reflect the dynamic nature of Paf1C association dynamics and H2Bub turn-over during transcription. However, we cannot exclude that the lack of a complete correlation between the presence of Paf1C and the H2Bub mark observed in some of these genes may be due to technical limitations linked to stringent peak calling for detecting them or to Paf1C-independent H2Bub dynamics.

The H2Bub mark preferentially accumulates in longer genes in *Arabidopsis* (Fig. S3D) (Roudier et al., 2011). In addition, in mouse muscle myoblast cells loss of Paf1C reduces H2Bub mark over the 5’ of long genes, while the entire gene body is affected in medium and small size genes (Hou et al., 2019). On this basis, we decided to determine whether the requirement of Paf1C for histone H2B monoubiquitination is affected by gene size in plants. We ranked the H2Bub- and Paf1C-marked genes by gene size and compared the H2Bub signal in wild-type and *elf7-3* seedlings. This showed that the impact of Paf1C on H2Bub level is independent of gene size in *Arabidopsis* (Fig. S4).

Studies in yeast and mammals provided molecular insights into the functional relationship between Paf1C and histone H2B monoubiquitination by the E2 conjugating enzyme Rad6 and the E3 ligase Bre1 in yeast and RNF20/RNF40 in mammals. In yeast, Paf1C promotes monoubiquitination through direct interaction with Rad6/Bre1 and does not appear to affect Rad6 recruitment to chromatin (Ng et al., 2003; Wood et al.,

2003; Kim and Roeder, 2009), whereas in mammals, RNF20/40 recruitment to chromatin is dependent on Paf1C (Wu et al., 2014). To investigate the situation in *Arabidopsis*, we examined whether the reduction in H2Bub mark in the *elf7-3* mutant was due to impaired recruitment to chromatin of HISTONE MONOUBIQUITINATION1 (HUB1) and HUB2, the two E3 ubiquitin ligases mediating H2Bub deposition in *Arabidopsis* (Liu et al., 2007). To this end, we fractionated cell extracts of the wild type and the *elf7-3* mutant and used a custom-made antibody to determine HUB1/2 levels (Fig. 3A) and their association with chromatin compared with cytoplasmic and nucleoplasmic fractions by immunoblotting. We observed a band corresponding to HUB1/2 in the chromatin fraction of wild-type cell extracts, and these levels were not affected by the *elf7-3* mutation (Fig. 3B). This result suggests that Paf1C promotes histone H2B monoubiquitination independently of HUB1/2 recruitment to chromatin in *Arabidopsis*.

**Figure 3.**
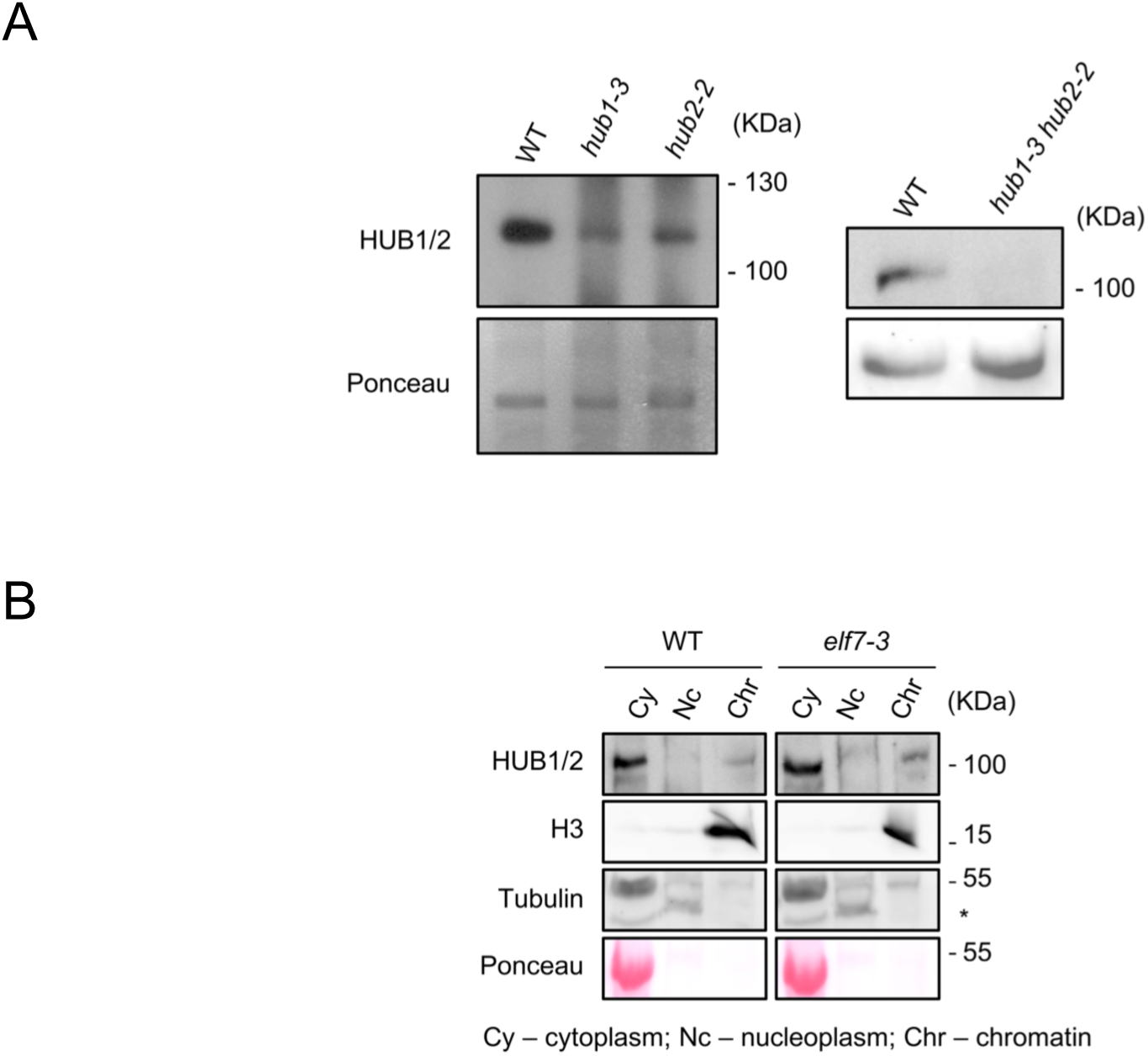
The recruitment of HUB1 and HUB2 to chromatin is independent of Paf1C. **A)** Immunoblot showing the specificity of the antiserum raised against HUB1 and HUB2. The bottom panels show the Ponceau staining. **B)** Detection of HUB1/HUB2 in cytoplasmic, nucleosplasmic, and chromatin fractions by Western blot analysis. Histone H3 was used as a chromatin marker and tubulin and Ponceau staining as cytoplasmic markers. The position of the molecular weight markers is shown on the right. The asterisk marks a non-specific band detected with the anti-tubulin antibody.

### Paf1C and histone H2B monoubiquitination regulate common processes

The functional relationship between Paf1C and H2Bub should manifest itself in common DEGs in mutants defective for Paf1C or for the machinery that monoubiquitinates histone H2B. We compared the transcriptomes of *elf7-3* and *hub1-4* mutant (Liu et al., 2007) seedlings grown side-by-side. We identified 413 DEGs (*p* adj. < 0.05) in *hub1-4* seedlings, a reduced number compared to *elf7-3* (Fig. 4, A and B). More than half of DEGs in *hub1-4* were also misregulated in *elf7-3* seedlings (Fig. 4B) and, importantly, this subset showed the same trend in gene expression change, in line with the functional link between Paf1C and H2Bub (Fig. 4, C and D). The majority of DEGs in *hub1-4* and those common to both mutants were occupied by Paf1C (Fig. S5), suggesting direct functional links between Paf1C and H2Bub in gene expression.

**Figure 4.**
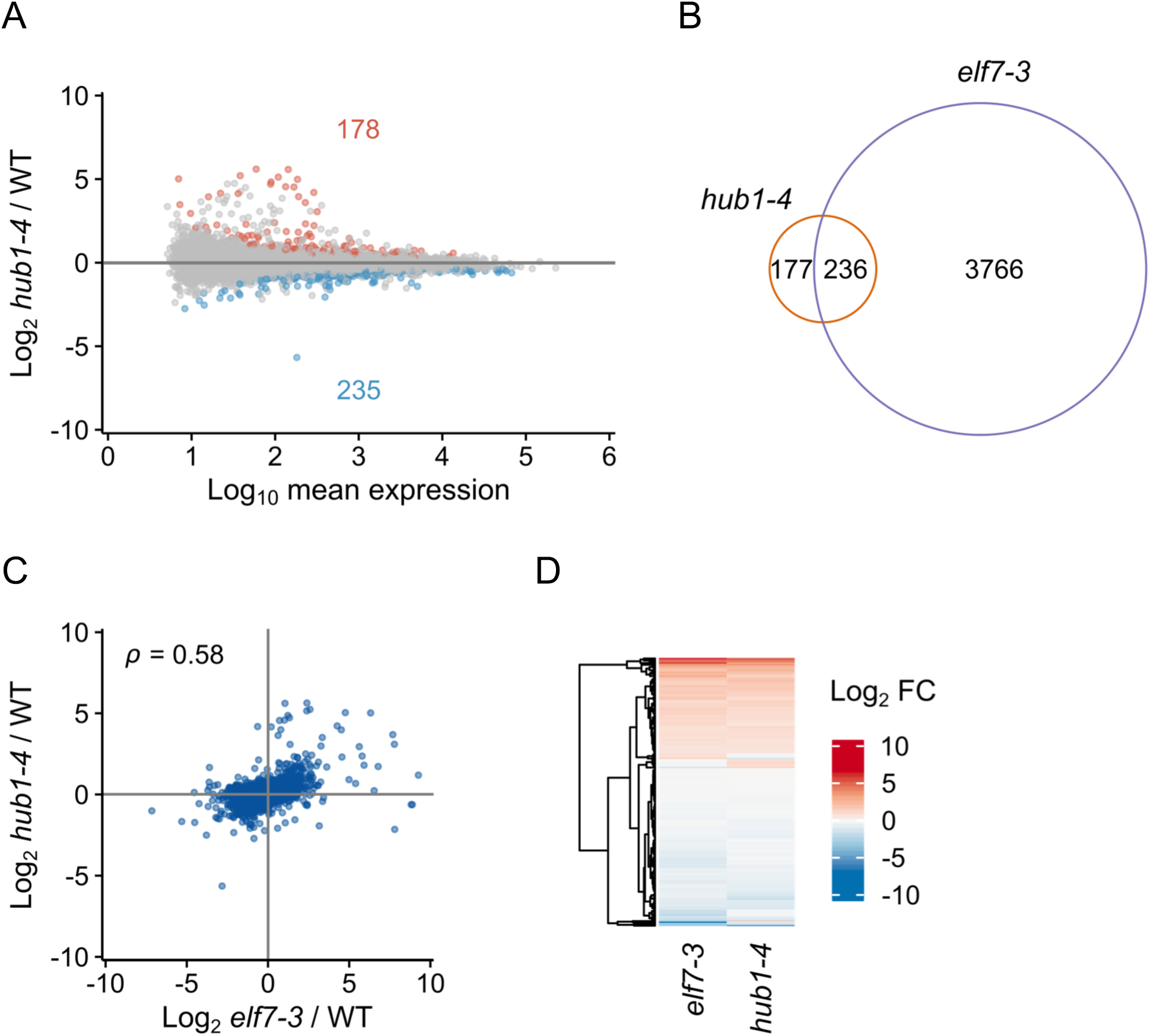
HUB1 and ELF7 regulate a common set of genes. **A)** MA plot showing DEGs (*p* adj. < 0.05) between *hub1-4* and wild-type seedlings. **B)** Comparison of DEGs between *hub1-4* and *elf7-3* seedlings. **C)** Scatter plot showing the correlation of common DEGs between *hub1-4* and *elf7-3* mutant seedlings. **D)** The heatmap shows the behavior of common DEGs between *hub1-4* and *elf7-3* mutant seedlings.

### *elf7* mutation causes RNAPII stalling at the 5’ end of genes

We next examined the effect of impaired Paf1C on RNAPII occupancy. ChIP-seq profiling of the NRPB1 subunit, which indiscriminately captures all active RNAPII states, detected peaks at 18,339 genes, half of which were also occupied by Paf1C. In wild-type plants, RNAPII distribution showed a typical pattern with peaks immediately after the transcription start site (TSS) and at the TES and lower levels throughout the gene body, where RNAPII processivity is high and residence time is low (Figs. 1B, 2B, and 5A). The peak after the TSS reflected RNAPII stalled at the +1 nucleosome (Fig. 5B), as previously reported (Kindgren et al., 2019). Importantly, we observed a redistribution of RNAPII signal in the *elf7-3* mutant compared with wild type plants, with reduced accumulation over gene bodies and TES and increased accumulation across the TSS region (Fig. 5, A and C). Overall, these defects most likely reflect a more frequent stalling of RNAPII in *elf7-3* plants.

**Figure 5.**
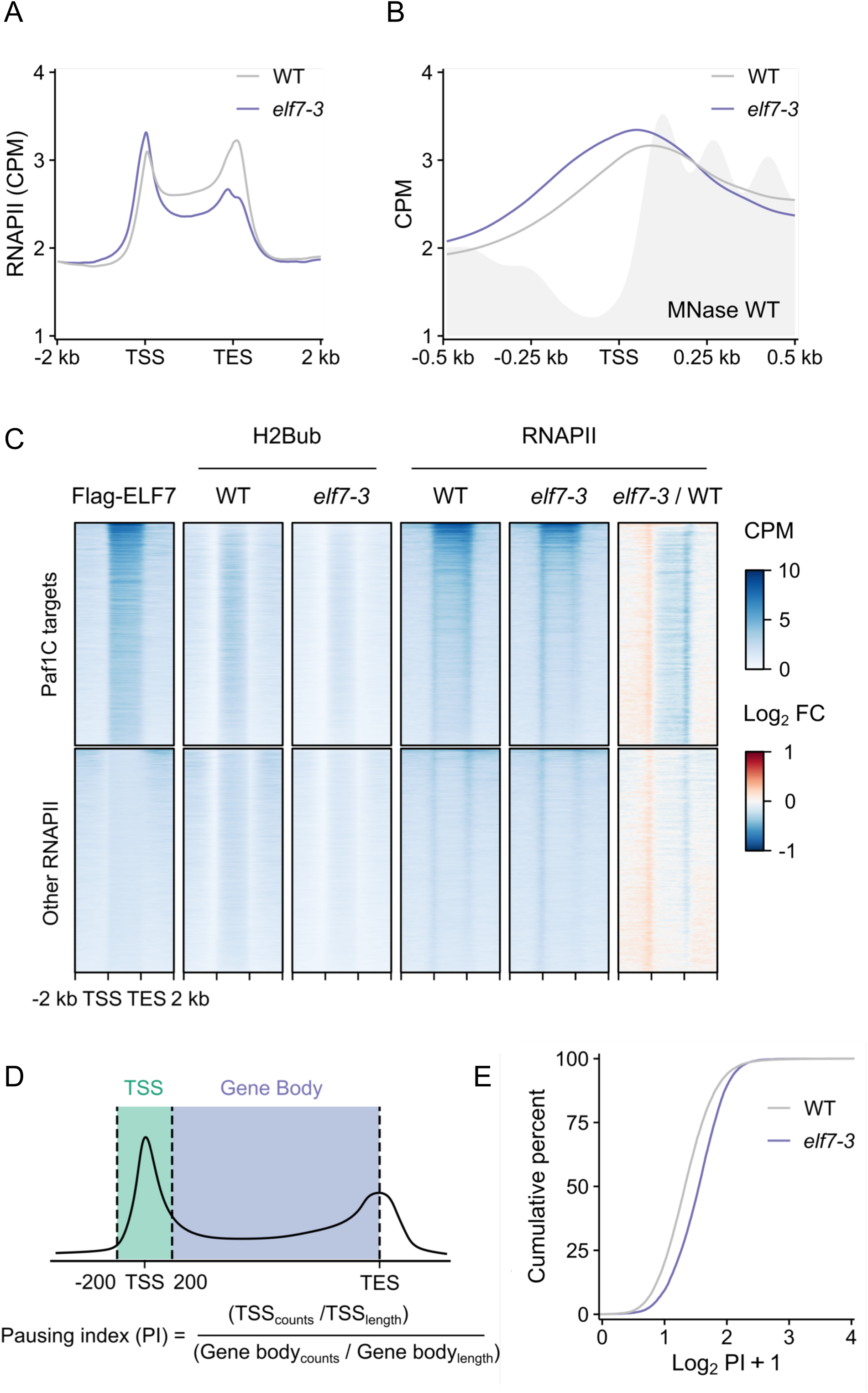
RNAPII genomic distribution is affected in *elf7-3* seedlings. **A)** The metagene analysis shows the average distribution of all RNAPII-occupied genes in wild type and in *elf7-3* mutants. **B)** Average distributions of RNAPII in wild type and *elf7-3* seedlings centered on the TSS, and compared with the nucleosome position previously determined by MNase-seq (Diego-Martín et al., 2022). **C)** The heatmap shows the effect of the *elf7-3* mutation on the distribution of RNAPII and H2Bub in genes occupied by Paf1C (Paf1C targets) and in genes in which we could not detect binding of Paf1C (Other RNAPII). **D)** Scheme depicting the calculation used to determine the pausing index (PI). **E)** Representation of the PI in wild type and *elf7-3* mutant. The higher the PI, the greater the degree of paused RNAPII.

Several lines of evidence pointed to a direct contribution of Paf1C to RNAPII redistribution. Reduction of RNAPII occupancy at gene bodies and TES in *elf7-3* seedlings was observed preferentially for Paf1C target genes (Fig. 5C). This suggests that Paf1C contributes locally to transcription elongation. Indeed, Paf1C target genes were enriched in histone marks associated with active transcription (Fig. 5C and Fig. S2A). However, the increased accumulation of RNAPII at TSS observed in *elf7-3* seedlings also occurred in genes where no Paf1C peak was detected (Fig. 5C). This result suggests that the effect of the *elf7-3* mutation on stalled RNAPII may result from direct and indirect consequences of defective Paf1C, such as reduced H2B monoubiquitination (Fig. 5C). Interestingly, the accumulation of stalled RNAPII in *elf7-3* mutant plants was shifted toward the 5’ end of the genes compared with wild type (Fig. 5B). However, it is unclear whether this shift is due to the +1 nucleosome moving toward the 5’ end of the genes in the mutant.

Finally, the possibility that Paf1C is involved in transcription elongation was further supported by comparing the RNAPII pausing index of wild-type and *elf7-3* mutant plants. Similar to previous studies (Wang et al., 2023a), we calculated the pausing index as the ratio of RNAPII occupancy across the TSS to RNAPII occupancy in the gene body (Fig. 5D). As shown in Fig. 5E, the RNAPII pausing index increased in plants of the Paf1C mutant. This is consistent with the idea that Paf1C contributes to the productive elongation of RNAPII.

## DISCUSSION

In this study, we provide genome-wide evidence for the function of Paf1C as a transcriptional elongation factor in plants. First, the Paf1C subunits ELF7 and VIP3 are exclusively distributed in gene bodies and their accumulation positively correlates with that of RNAPII and gene expression, suggesting that the complex actively promotes gene transcription (Fig. 1, B-D and Fig. S2). Second, monoubiquitination of histone H2B, a chromatin mark that is deposited co-transcriptionally, is severely reduced in virtually all genes in Paf1C-defective plants, a defect that is particularly pronounced in Paf1C-targeted genes (Figs. 2 and 4C). Third, RNAPII appears to stall more frequently at the 5’ end of genes, and the overall gene body occupancy is reduced in Paf1C-targeted genes in *elf7-3* mutant plants (Fig. 4A-C). This likely reflects the inability of RNAPII to transition to the productive elongation stage. The latter is consistent with previous results in *Arabidopsis*, which demonstrate that (i) the phosphorylated form of RNAPII at serine 5 (Ser5) of the C-terminal domain of NRPB1 (RNAPII-Ser5), marking engaged RNAPII, accumulates in the *elf7-3* mutant around the TSS (Wang et al., 2023b); and (ii) the occupancy at gene bodies of Ser2 phosphorylated RNAPII, the elongating form, is reduced in the *elf7-3* mutant (Obermeyer et al., 2022).

Despite this evidence for a role of Paf1C in transcription elongation, we still lack information on the mechanism of action of this complex in plants. In this regard, a comparison of the genomic distribution of Paf1C and RNAPII in different model organisms could provide relevant information. The genomic profile of the *Arabidopsis* Paf1C subunits ELF7 and VIP3 strongly resembles that of the Paf1 subunit in yeast, as both are mainly localized within the gene bodies (Fig. 1, B and C) (Mayer et al., 2010; Wang et al., 2023b). It is noteworthy, however, that while RNAPII, much like Paf1C, is found within gene bodies, their distributions do not entirely overlap, as RNAPII exhibits heightened enrichment at the TSS and TES (Figs. 1B and 5B). In mammals, the distribution of Paf1C parallels the distribution of RNAPII, as both are enriched across the TSS and after the TES (Rahl et al., 2010; Chen et al., 2015; Yu et al., 2015). In mammals, their similar distributions therefore reflect an association of Paf1C with RNAPII, with Paf1C involved in the regulation of RNAPII pausing immediately after the TSS (Chen et al., 2015; Yu et al., 2015) and contributing to mRNA polyadenylation (Nagaike et al., 2011). However, in *Arabidopsis*, the genes are generally much shorter than in mammals. Consistent with the positive correlation observed in mammals between the distribution of Paf1C and RNAPII, the accumulation of Paf1C at gene bodies in *Arabidopsis* may be important for productive transcription elongation, as shown in Fig. 5C. Consistent with this possibility, RNAPII accumulates near the TSS of genes and is locally shifted toward 5’, whereas the RNAPII pausing index is increased in the *elf7* mutant compared with wild-type plants.

How does Paf1C contribute to transcriptional elongation? The association of Paf1C with RNAPII could facilitate the passage of the transcriptional machinery through the nucleosomes through several mechanisms. Structural studies in mammals have shown that Paf1C position in the RNAPII transcriptional complex should enable it to promote methylation of histone H3K4 in downstream nucleosomes -a modification known to facilitate transcription (Vos et al., 2018). We hypothesize that Paf1C would be recruited to RNAPII positioning itself within the complex similarly to its mammalian counterpart. Subsequently, Paf1C would promote post-translational modifications of histones such as monoubiquitination of H2B in nucleosomes downstream of the +1, thereby facilitating the passage of RNAPII thoughout the whole gene body where we found both Paf1C and H2Bub to frequently accumulate. The direct effect of Paf1C on local HUB1/2 activity is supported by several observations in this study, including the frequent co-occupancy of genes by Paf1C and RNAPII, with high levels of H2Bub in this context, the association of HUB1/2 with chromatin even in the absence of Paf1C, and the previously described physical interaction between the Paf1C subunit VIP5 and the *Arabidopsis* homologs of the Rad6 E2 ubiquitin-conjugating enzyme (Li et al., 2023). Nevertheless, the reduced H2Bub deposition in *elf7* mutant plants could in some cases result from indirect effects of Paf1C deficiency, such as impaired RNAPII elongation.

## MATERIALS AND METHODS

### Plant material

The *Arabidopsis* lines used in this work have been previously described: *elf7-3* (He et al., 2004), *hub1-4* and *hub1-3 hub2-2* (Liu et al., 2007), *pELF7:Flag-ELF7 elf7-3* (Cao et al., 2015), and *35S:GFP-VIP3* (Dorcey et al., 2012).

### Growth conditions and treatments

All seeds were surface sterilized and sown on half-strength MS (Duchefa) plates containing 1% (w/v) sucrose, 8 g/L agar (pH 5.7). Seedlings were grown at 22°C under continuous light (50-60 μmol m^−2^ s^−1^) (standard conditions).

### RNA-seq experiments

RNA-seq with three independent biological replicates for each genotype were performed. Seedlings were grown as described above. 7-day-old wild type, *elf7-3*, and *hub1-4* seedlings were collected. Total RNA was extracted using an RNeasy Plant Mini Kit (Qiagen) according to the manufactureŕs instructions. RNA concentration and integrity were measured in an RNA Nanochip (Bioanalyzer, Agilent Technologies 2100) from IBMCP Genomics Service. Library preparation and sequencing were performed by the Genomics Service of the University of Valencia.

### RNA-seq analysis

RNA-seq read quality was first evaluated using FastQC v0.11.9. Low-quality bases and Illumina adapters were trimmed using cutadapt v4.2 with options “-q 15,10 -a AGATCGGAAGAGCACACGTCTGAACTCCAGTCA -m 20 --trim-n”. Clean reads were then aligned to the TAIR10 reference genome using STAR v2.7.10b (Dobin et al., 2013) with the default parameters. Read counts were obtained using the featureCounts command from subread v2.0.3 (Liao et al., 2014) with the default parameters and providing the Araport11 annotation. Differential analyses were performed using DESeq2 v1.38 (Love et al., 2014). Genes showing an adjusted p-value inferior to 0.05 were considered as differentially expressed.

### ChIP experiments

H2B and H2Bub ChIP-Rx were performed in parallel with the same two biological replicates of 7-day-old wild-type and *elf7-3* seedlings grown under standard conditions. As previously described (Nassrallah et al., 2018), all samples were spiked-in with *Drosophila* chromatin prior to IP. The amount of exogenous DNA was subsequently determined in all input and IP samples and considered as a reference to avoid the effects of technical variation. Furthermore, we determined the H2Bub peaks after normalization to histone H2B occupancy determined in the same samples by ChIP-Rx. Specifically, for each biological replicate, two IPs were performed with an anti-H2Bub antibody (MM -0029-P, Medimabs) and one with an anti-H2B (ab1790, Abcam). For each IP, 100 µg of *Arabidopsis* chromatin mixed with 3 µg of *Drosophila* chromatin was used. DNA eluted from the two technical replicates of the H2Bub IP was pooled prior to library preparation. Library preparation and sequencing was performed by the CRG Genomics Core Facility (Barcelona, Spain). Flag-ELF7 and GFP-VIP3 ChIP-seqs were performed using 7-day-old seedlings grown under standard conditions and an anti-Flag M2 antibody (F1804, Sigma) and an anti-GFP antibody (ab290, abcam), respectively. We used double in vitro cross-linking with 1.5 mM ethylene glycol bis (succinimidyl succinate) for 20 minutes followed by 1% formaldehyde for 10 minutes at room temperature. Library preparation and sequencing were performed by the CRG Genomics Core Facility (Barcelona, Spain). ChIP-seq for RNAPII was performed using an anti-RPB1 antibody (clone 4H8, Active Motif) using 7-day-old wild-type and *elf7-3* seedlings grown under standard conditions. Library preparation and sequencing were performed by the Epigenomics platform at IPS2 (Paris, France).

### ChIP-seq analysis

The quality of ChIP-seq reads was first assessed using FastQC v0.11.9. Low-quality bases and Illumina adapters were trimmed using cutadapt v4.2 with options “-q 15,10 -a AGATCGGAAGAGCACACGTCTGAACTCCAGTCA -m 20”. Clean reads were then aligned to the TAIR10 reference genome using bowtie2 v2.5.1 (Langmead and Salzberg, 2012) with the default parameters. Alignments were sorted by coordinate using the samtools sort command from samtools v1.17 (Li et al., 2009). Duplicate reads were then marked using the sambamba markdup command from sambamba v1.0 (Tarasov et al., 2015) and uniquely mapped reads were retained using the samtools view command with options “-F 4 -F 1024 -q 5”. Coverage files in bedGraph format were obtained using the genomeCoverageBed command from bedtools v2.31.0 (Quinlan, 2014) with options “-bga -fs 200”, and then these were converted to bigwig using the bedGraphToBigWig command. Bigwigs were normalized to Counts Per Million (CPM) using the wiggletools scale command from wiggletools v1.2 (Zerbino et al., 2014). Mean coverages of biological replicates from each condition were obtained using the wiggletools mean command. Log2 ratios between treatments and controls were obtained using the bigwigCompare command from deeptools v3.5.1 (Ramírez et al., 2014).

Peak calling was performed over uniquely mapped reads using macs2 callpeak from macs2 v2.2.7.1 (Zhang et al., 2008) with the following parameters: “--keep-dup all -- broad --broad-cutoff 0.05 -q 0.01 --nomodel --extsize 200”. Peaks from biological replicates were pooled and merged into a consensus set using the bedtools merge command from bedtools v2.31.0 (Quinlan, 2014). In experiments that included various treatments (ChIP-seq of H2Bub and RNAPII), the consensus peaks were obtained by merging the peaks from each treatment using the same procedure as for the biological replicates. Consensus peaks were annotated to their closest gene using the bedtools closest command. To determine which genes could be considered as targets, the following filters were applied: the peak must overlap more than half of the gene body, or the gene body must overlap more than half of the peak, or the peak center must be less than 200 pb away from the TSS.

### ChIP-Rx analysis

H2Bub ChIP-Rx reads were first processed the same way as ChIP-seq reads. Clean reads were aligned against a chimeric genome that contained both *Arabidopsis* (TAIR10) and *Drosophila* (dm6) chromosomes. Alignments were sorted and duplicate-marked as in ChIP-seq, and then reads that mapped to TAIR10 or to dm6 were separated. TAIR10 alignments were filtered and used to create coverage bigwigs and for peak calling as previously described. Rx normalization factors were calculated as described in Nassrallah et al., (2018). Briefly, the normalization factor α for each ChIP sample was obtained using the following formula:

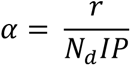

where N_d_ is the number of mapped reads to *Drosophila* dm6 in millions and r is the percentage of dm6 reads in the corresponding input sample. Hence, r was calculated as:

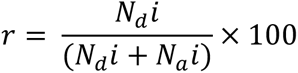

where N_d_i and N_a_i are the number of reads mapped to dm6 and TAIR10.

To test differential ubiquitination, read counts over the consensus H2Bub peaks were first obtained using the multiBamSummay command from deeptools v3.5.1 (Ramírez et al., 2014) with options “--extendReads 200 --minMappingQuality 5”. Gene level counts were obtained by summing counts from peaks that were annotated to the same gene. Differential analyses were performed using DESeq2 v1.38.0 (Love et al., 2014). In these analyses, the Rx normalization factors were provided as size factors. Genes showing an adjusted p-value inferior to 0.05 were considered as differentially ubiquitinated.

### Pausing index

To estimate RNAPII pausing, the pausing index formula was employed (Wang et al., 2023a):

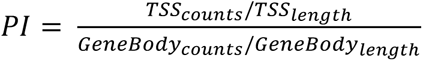

The TSS was defined as a 400 pb region centered in the annotated TSS, whereas the gene body spanned from the TSS region to the annotated TES. For this reason, only RNAPII genes that were longer than 200 pb were included. Bedfiles of TSSs and gene bodies were created using the slopBed command from bedtools v2.31.0 (Quinlan, 2014) and the TAIR10 reference annotation. Counts over these regions were then obtained using the multiBamSummary command from deeptools v3.5.1 (Ramírez et al., 2014).

### Re-analysis of existing datasets

ChIP-seq reads of H3, H3K4me3, H3K36me3, H3K27me3, and H2A.Z (GSE231408; samples GSM7075958, GSM7075959, GSM7075961, GSM7075962, GSM7075963, GSM7075964, GSM7075985, GSM7075986 and GSM7075990) (Jamge et al., 2023) were analysed as described in ChIP-seq analysis. MNase-seq reads of wild type Col-0 (GSE205110; samples GSM6205325 and GSM6205326) (Diego-Martín et al., 2022) were downloaded, processed and aligned as described in ChIP-seq analysis. Coverage bigwigs were obtained using DANPOS v2.2.2 (Chen et al., 2013) and the wigToBigWig command.

### Data representations

Graphs were plotted in R 4.1.2 using ggplot2, eulerr and ComplexHeatmap libraries (Gu et al., 2016). Other heatmaps and metaplots were obtained using the computeMatrix, plotHeatmap and plotProfile commands from deeptools v3.5 (Ramírez et al., 2014).

### Subcellular fractionation

Cytoplasmic, nucleoplasmic, and chromatin-associated fractions were obtained following a described protocol (Liu et al., 2018), with some minor modifications. Briefly, approximately 1.5 g of 7-day-old seedlings were ground in liquid nitrogen and subsequently homogenized in 3-4 mL of Honda buffer (0.44 M Sucrose, 20 mM HEPES KOH pH 7.4, 2.5% Percoll, 5% Dextran T40, 10 mM MgCl_2_, 0.5% Triton X-100, 5mM DTT, 1 mM PMSF, and 1x protease inhibitor cocktail [cOmplete, EDTA-free; Roche]). The resulting homogenate was filtered through two layers of Miracloth, and the flow-through was then subjected to centrifugation at 2,400 *g* for 10 minutes at 4°C. The resulting supernatant (1 mL) was further centrifuged at 10,000g for 10 minutes at 4°C, and this supernatant was collected as the cytoplasmic fraction. The pellet was resuspended in 1 mL of Honda buffer and centrifuged at 1,800 *g* for 5 minutes at 4°C to concentrate nuclei. This pellet was subsequently washed 4-6 times with Honda buffer and rinsed with PBS buffer (137 mM NaCl, 2.7 mM KCl, 10 mM Na_2_HPO_4_, and 2 mM KH_2_PO_4_) containing 1 mM EDTA. The resulting pellet was resuspended in 150 µL of cold glycerol buffer (20 mM Tris-HCl pH 7.9, 50% glycerol, 75 mM NaCl, 0.5 mM EDTA, 0.85 mM DTT, 0.125 mM PMSF, and 1x protease inhibitor cocktail [cOmplete, EDTA-free; Roche]) and gently vortexed twice after adding 150 µL of cold nuclei lysis buffer (10 mM HEPES KOH pH 7.4, 7.5 mM MgCl_2_, 0.2 mM EDTA, 0.3 M NaCl, 1 M urea, 1% NP-40, 1 mM DTT, 0.5 mM PMSF, 10 mM β-mercaptoethanol, and 1x protease inhibitor cocktail [cOmplete, EDTA-free; Roche]). The mixture was then incubated for 2 minutes on ice and centrifuged at 14,000 rpm for 2 minutes at 4°C. The resulting supernatant was collected as the nucleoplasmic fraction. The chromatin-associated pellet was rinsed with PBS buffer containing 1 mM EDTA and then resuspended in 150 µL of cold glycerol buffer plus 150 µL of cold nuclei lysis buffer. Protein concentrations were determined using the Pierce 660 nm Protein Assay following the manufacturer’s instructions. The various fractions were subsequently analyzed by immunoblot.

### Immunoblot analysis

Total protein extracts and subcellular fractionation samples were separated with SDS-PAGE, transferred to PVDF membranes, and immunolabeled with commercial antibodies against histone H3 (ab1791, Abcam) and against tubulin (62204, Invitrogen). Chemiluminescence was detected using the Supersignal west FEMTO substrate with maximum sensitivity (Thermo-Fisher Scientific), and protein bands were detected and quantified using the LAS-3000 Imaging System (Fujifilm) and ImageJ software, respectively. The custom-made rabbit anti-HUB1 antibody was generated for F. Barneche by SDIX (Newark, USA) using the following immunogen sequence MQDTLLIDKYIMDKDIQQGSAYASFLSKKSSRIEDQLRFCTDQFQKLAEDKYQKSVSLENL QKKRADIGNGLEQARSRLEESHSKVEQSRLDYGALELEL.

## ACCESSION NUMBERS

The sequencing data generated in this work have been deposited in the GEO public functional genomics data repository under the accession number GSE244850.

## SUPPLEMENTAL DATA

Supplemental document 1: Supplemental Figures S1-S5.

Supplemental document 2: Supplemental Table.

## ACKNOWLEDGEMENTS AND FUNDING

We thank Christian S. Hardtke (University of Lausanne, Switzerland) and Ligeng Ma (Capital Normal University, China) for sharing seeds of the *35S:GFP-VIP3* line and of *elf7-3* and *pELF7:Flag-ELF7 elf7-3*, respectively. N.B-T was supported by a predoctoral contract from the MINECO [BES-2014-068868] and an EMBO Short-Term fellowship (STF n°8047) to visit the Barneche/Bourbousse laboratory. J.P-A was supported by a predoctoral contract from the Generalitat Valenciana (CIACIF/2021/432). Research in the IBMCP laboratories was funded by grants BIO2013-43184-P (to M.A.B. and D.A.) and BIO2016-79133-P (to D.A.) from the Spanish Ministry of Economy and Innovation, as well as grants PID2019-109925GB-I00 (to D.A.) and RYC2018-024108-I and PID2019-108577GA-I00 (to J.G-B) from MCIN/AEI/10.13039/501100011033. Research in the Barneche/Bourbousse laboratory was funded by Agence Nationale de la Recherche grants ChromaLight (ANR-18-CE13-0004-01), Epilinks (ANR-22-CE20-0001) and PlastoNuc (ANR-20-CE13-0028). Research in the Benhamed laboratory was funded by the European Research Council ERC (Project 101044399-3Dwheat), Agence National de la Recherche ANR (ANR-21-CE20-0036-4D Heat Tomato) and by the Institut Universitaire de France (IUF).

## AUTHOR CONTRIBUTIONS

N.B-T., J.P-A., C.B., M.B., F.B., M.A.B., J.G-B., and D.A. designed the research; N.B-T., J.P-A., C.B., D.L., O.A., M.B., F.B., J.G-B., and D.A. performed research; N.B-T., J.P-A., C.B., D.L., O.A., M.B., F.B., M.A.B., J.G-B., and D.A. analyzed data; D.A. wrote the paper.

