## Supplemental Figures and Figure Legends for "Genome-wide evidence of the role of Paf1C in transcription elongation and histone H2B monoubiquitination in *Arabidopsis*"

**Supplemental Figure Legends**

**Figure S1.** The Paf1C subunits ELF7 and VIP3 bind to common targets. **A, B)** Scatter plots showing the correlation between replicates of Flag-ELF7 (A) and GFP-VIP3 (B) ChIP-seq experiments. **C)** Examples of Flag-ELF7 and GFP-VIP3 targets extracted from the IGV.

**Figure S2.** Binding of ELF7 and VIP3 correlates with epigenetic marks associated to active gene expression. **A)** Heatmaps showing the signal of ELF7, VIP3, RNAPII, and several epigenetic marks and histone variants. Genes are ranked by their decreasing ELF7 signal. **B)** Heatmaps comparing the results of the ChIP-seq analisis of Flag-ELF7 done in this work and that previously described (Wang et al., 2023). **C)** Average plot of Flag-ELF7, GFP-VIP3, and RNAPII signal in DEGs categories "Paf1C targets", "Other expressed", and "Not expressed" found in *elf7-3* seedlings.

**Figure S3.** Dependent and independent effects of Paf1C on H2Bub marking. **A)** Venn diagram showing the overlap of genes occupied by Paf1C, RNAPII, and H2Bub. **B)** Examples of H2Bub occupancy extracted from the IGV showing the effect of the *elf7-3* mutation. **C)** Examples of H2Bub occupancy extracted from the IGV showing the ocuppancy of H2Bub and the Paf1C subunits ELF7 and VIP3. **D)** Violin plots showing the average gene size between those occupied by Paf1C and H2Bub and H2Bub or Paf1C alone.

**Figure S4.** The effect of Paf1C on H2Bub is independent on gene size. The heatmaps show the H2Bub signal in wild type and *elf7-3* seedlings ranked by increasing gene size.

**Figure S5.** Venn diagram showing the overlap between common DEGs of *hub1-4* and *elf7-3* mutants and Paf1C targets.

### Figure S1

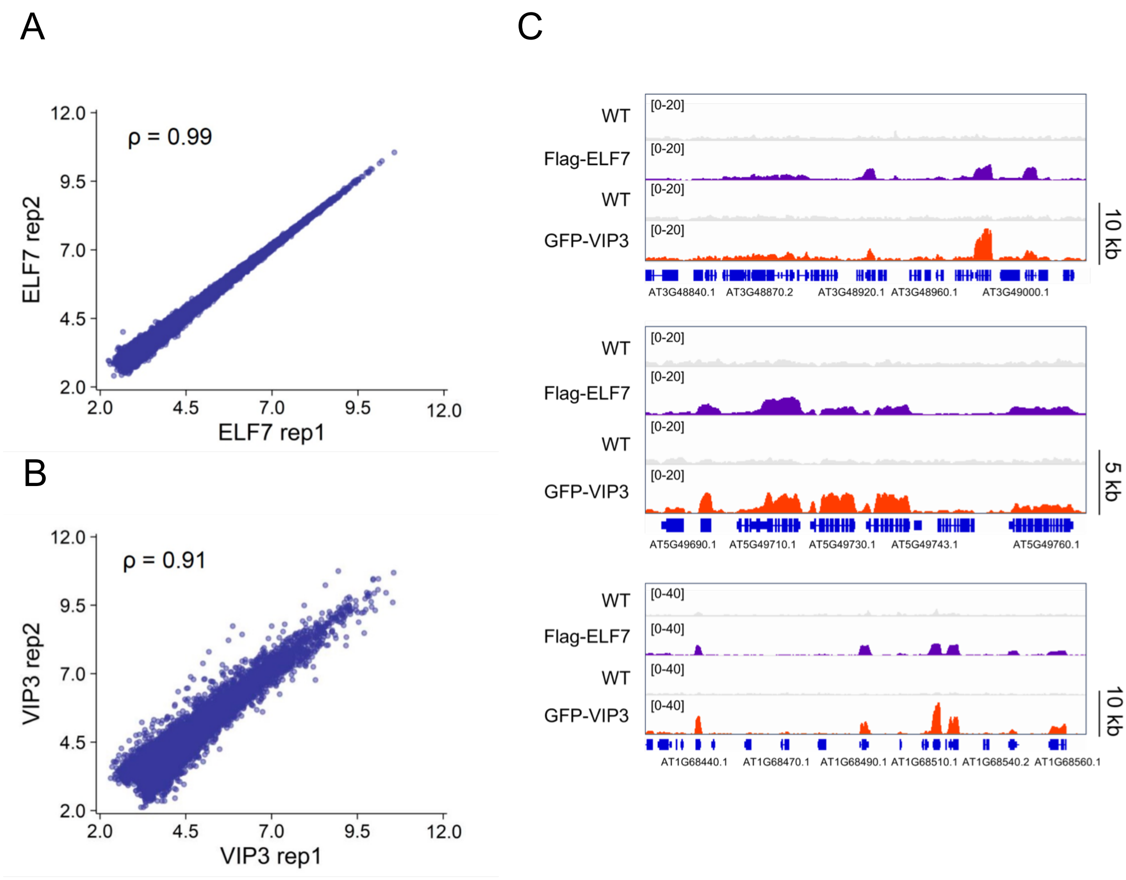
A
ρ = 0.99
ELF7 rep2
ELF7 rep1
12.0
9.5
7.0
4.5
2.0
B
ρ = 0.91
VIP3 rep2
VIP3 rep1
C
WT
Flag-ELF7
WT
GFP-VIP3
[0-20]
[0-40]
10 kb
5 kb
AT3G48840.1
AT3G48870.2
AT3G48920.1
AT3G48960.1
AT3G49000.1
AT5G49690.1
AT5G49710.1
AT5G49730.1
AT5G49743.1
AT5G49760.1
AT1G68440.1
AT1G68470.1
AT1G68490.1
AT1G68510.1
AT1G68540.2
AT1G68560.1

### Figure S2

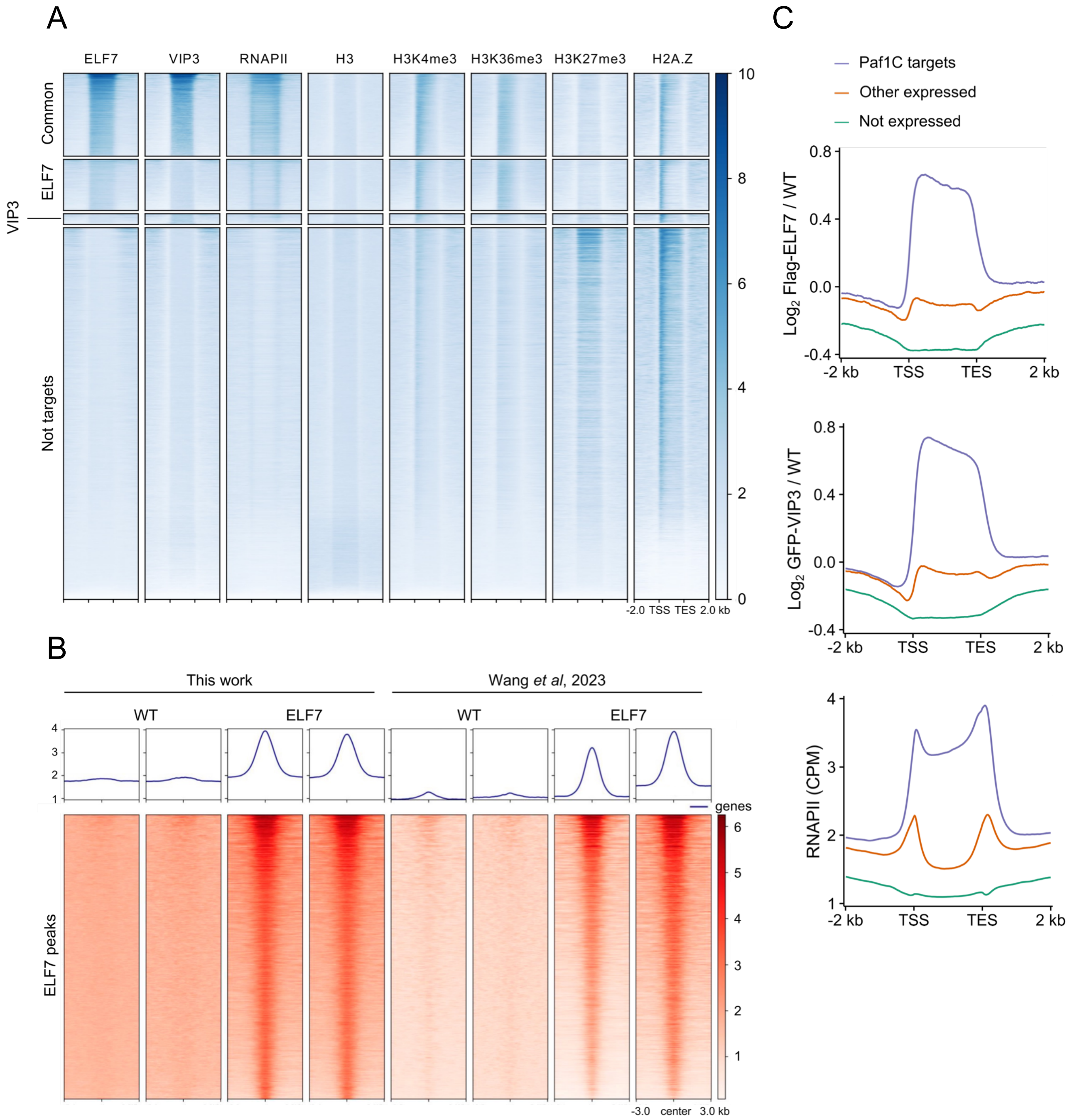

A
ELF7
VIP3
RNAPII
H3
H3K4me3
H3K36me3
H3K27me3
H2A.Z
Common
ELF7
VIP3
Not targets
10
8
6
4
2
0
-2.0 TSS TES 2.0 kb
B
This work
Wang et al, 2023
WT
ELF7
WT
ELF7
genes
ELF7 peaks
-3.0 center 3.0 kb
C
Paf1C targets
Other expressed
Not expressed
Log2 Flag-ELF7 / WT
Log2 GFP-VIP3 / WT
RNAPII (CPM)
-2 kb
TSS
TES
2 kb

### Figure S3

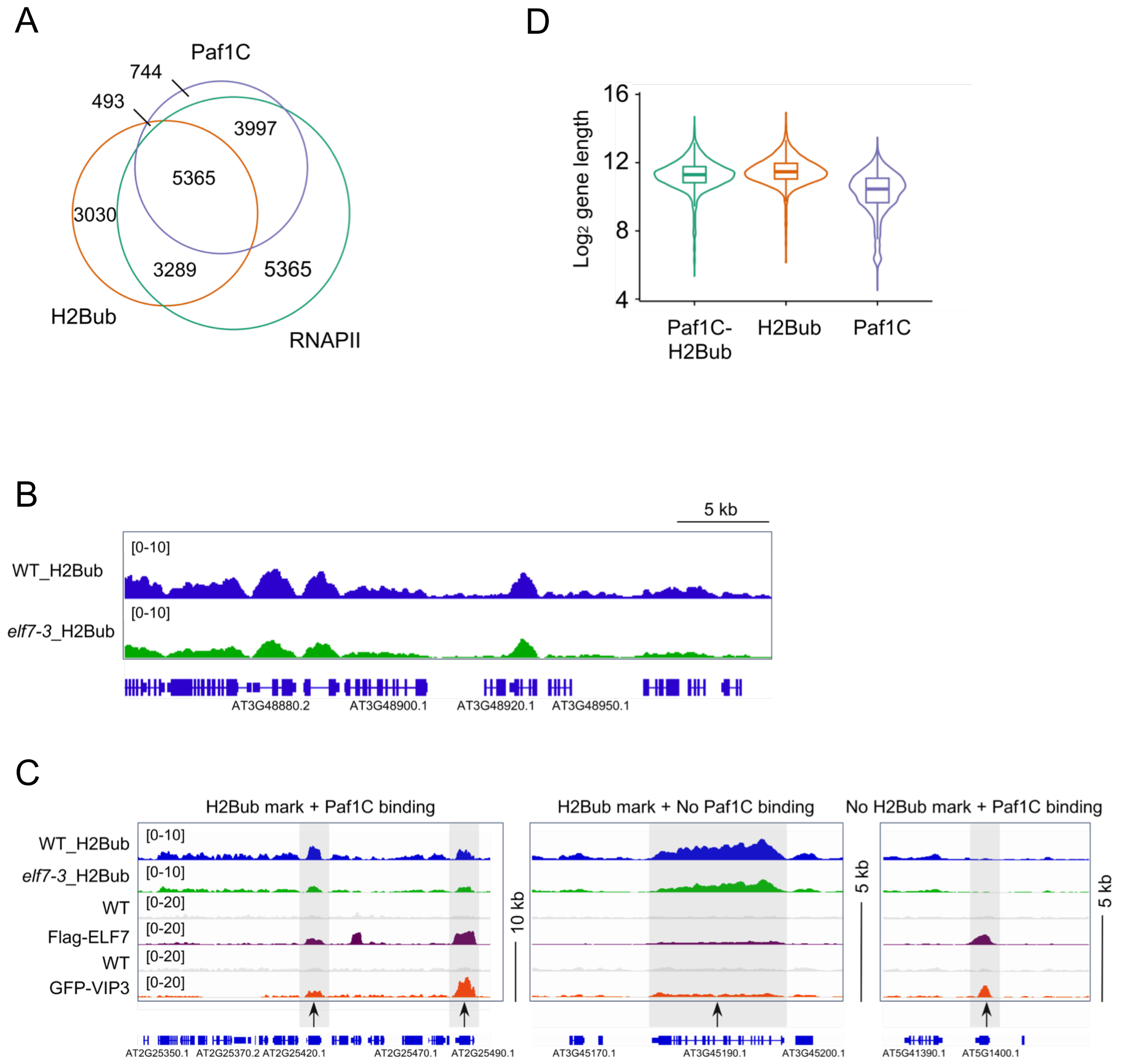
A
Paf1C
744
493
3997
5365
3030
3289
5365
H2Bub
RNAPII
D
16
12
8
4
Log2 gene length
Paf1C-
H2Bub
H2Bub
Paf1C
B
5 kb
WT_H2Bub
[0-10]
elf7-3_H2Bub
[0-10]
AT3G48880.2
AT3G48900.1
AT3G48920.1
AT3G48950.1
C
H2Bub mark + Paf1C binding
H2Bub mark + No Paf1C binding
No H2Bub mark + Paf1C binding
WT_H2Bub
elf7-3_H2Bub
WT
Flag-ELF7
WT
GFP-VIP3
[0-10]
[0-10]
[0-20]
[0-20]
[0-20]
[0-20]
10 kb
5 kb
5 kb
AT2G25350.1
AT2G25370.2
AT2G25420.1
AT2G25470.1
AT2G25490.1
AT3G45170.1
AT3G45190.1
AT3G45200.1
AT5G41390.1
AT5G1400.1

### Figure S4

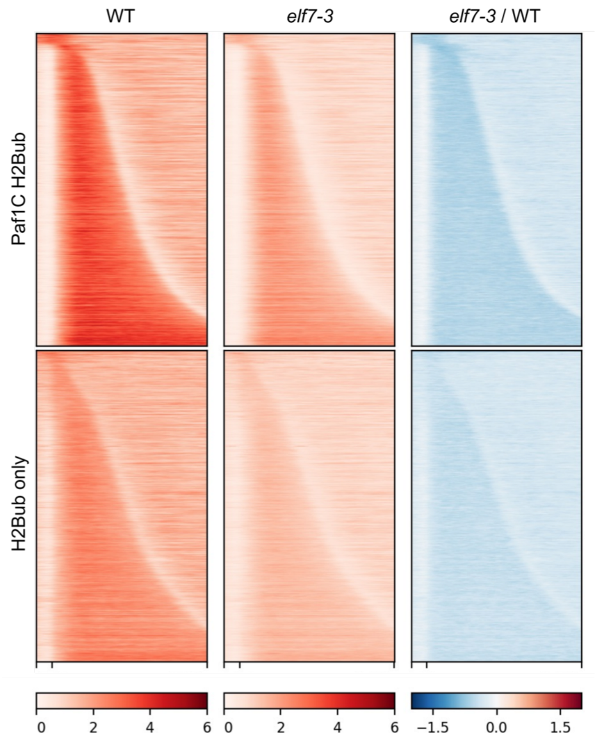

### Figure S5

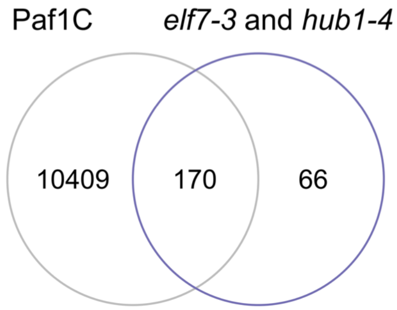
